# Deciphering the polycistronic nature of *Mycobacterium tuberculosis* lipoproteins of unknown functions

**DOI:** 10.1101/2023.07.16.549196

**Authors:** Blanco, Fabiana Bigi

**Affiliations:** Buenos Aires, Paraguay 2155 (C1121ABG), Buenos Aires, Argentina; Instituto de Biotecnología, CICVyA, Instituto Nacional de Tecnología Agropecuaria, Argentina (INTA), N. Repetto and de los Reseros, Hurlingham (1686), Buenos Aires, Argentina

## Abstract

*Mycobacterium tuberculosis*, the causative agent of human tuberculosis, encodes 116 lipoproteins, from which 48 have no demonstrated or predicted function. In this study, we demonstrated that six genes of these lipoproteins of unknown function are encoded in transcriptional units together with genes with experimentally verified functions or that encode conserved functional domains. We assigned predicted functions to LpqT (Rv1016c), LppE (1881c), LppO (Rv2290), LprE (Rv1252c) and LpqF (Rv3593).

## Introduction

*Mycobacterium tuberculosis* is a facultative intracellular bacterium that primarily infects the lungs and lymph nodes of humans, leading to tuberculosis. Although researchers have annotated the genome of *M. tuberculosis* in 1998 (Cole et al., 1998), the functions of many of its genes remain unknown. Among these genes, 116 are lipoprotein-encoding genes.

Regarding lipoproteins, these proteins are characterized for having a diacyl-glycerol moiety on the conserved cysteine present in the lipobox motif. Lipoproteins are mainly translocated across the plasma membrane by two transporters, the conserved Secretory (Sec) pathway and the Twin Arginine (Tat) system (Rezwan et al., 2007). In pathogenic mycobacteria, lipoproteins play several roles in different processes, including transport, cell wall biogenesis, defence and resistance, enzymatic and signalling processes and regulatory systems.

Based on the premise that genes encoded in the same operon are functionally connected, we focussed our analyses on the genes localised around lipoprotein-encoding genes of unknown functions. We demonstrated that five lipoprotein-encoding genes are transcribed together with genes that carry conserved domains or have an assigned function.

## Material and Methods

Template cDNA were obtained from *M. tuberculosis* H37Rv, CDC 1551 and *M. bovis* NCTC10772. Two μg of total RNA from exponential growth phase cultures of H37Rv, CDC1551 or NCTC10772 were purified using Trizol (Invitrogen) and treated with amplification grade DNAse (Invitrogen) to remove genomic DNA contamination. The cDNA synthesis was performed using 1 μg of RNA with M-MLV reverse transciptase (Promega) (RT+) or without this enzyme (mock reaction; without M-MLV enzyme) (RT-). The PCR reaction mixtures for the 11 intergenic regions were performed using specific primers (Table 1) in a final concentration of 300nM, dNTPs (Invitrogen), EasyTaq (Promega) and 1ul of template RT+ or RT-in a final volume of 20uL, following the manufacturer protocol. The amplification was performed with a standard program (95°C 5 min, followed by 35 cycles of 95°C 30 seg, 60°C 30 seg and 72°C 1 min and a final elongation step of 72°C for 10 min).

**Table 1:**
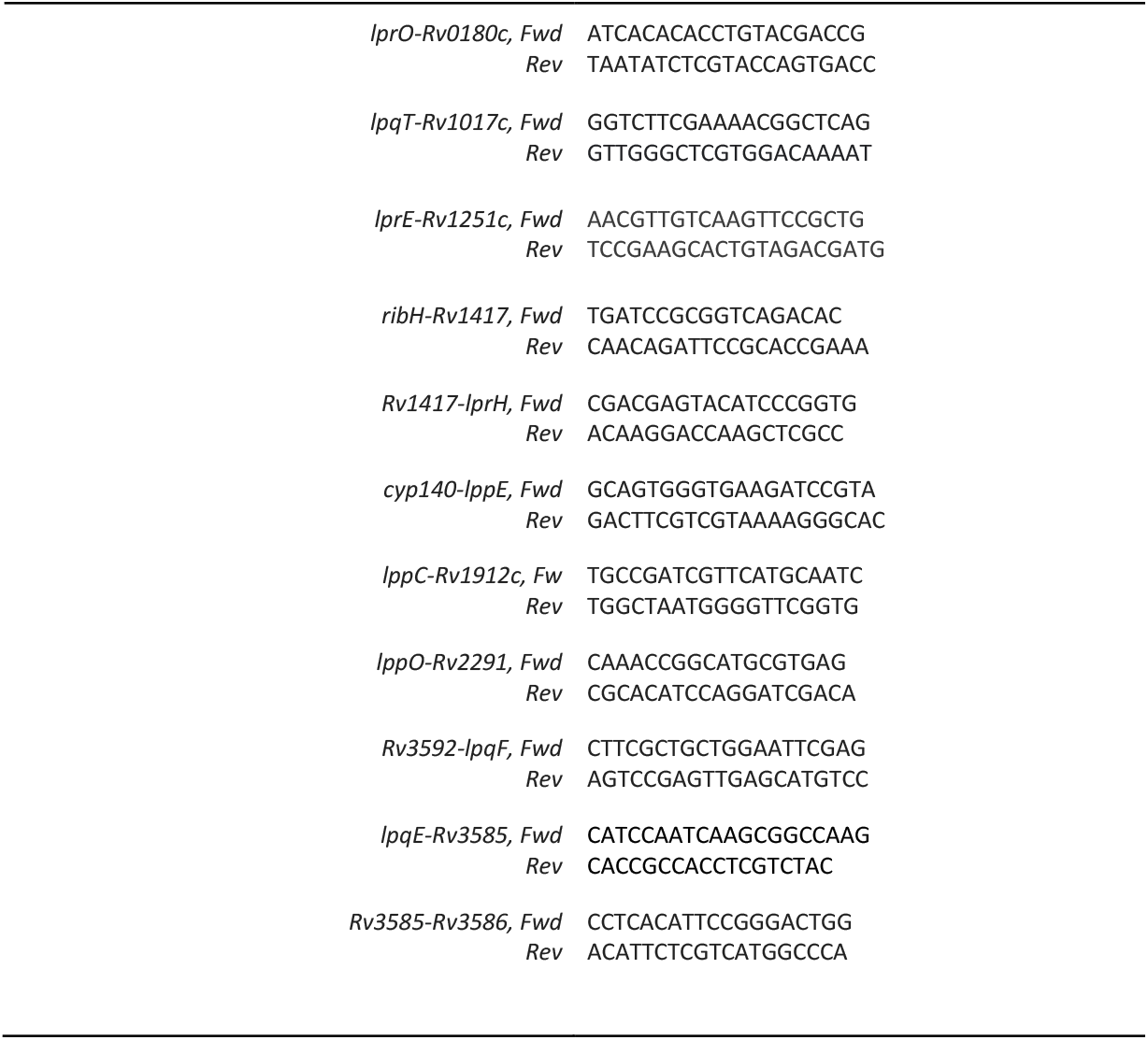
Primers used in RT-PCR.

## Results and discussion

The analysis using the Mycobrowser database (https://mycobrowser.epfl.ch/) repository yielded 45 lipoprotein-encoding genes that are genetically structured as possible operons. From these genes, at least ten encode lipoproteins of unknown functions. Total RNA from *M. tuberculosis* H37Rv, *M. tuberculosis* CDC 1551 and *Mycobacterium bovis* NCTC10772 were obtained to perform a transcriptional analysis from seven genes out of the ten identified lipoproteins of unknown functions. The RNA samples were retrotranscribed to cDNA and used as templates for PCR analysis. Pairs of primers were designed to amplify the lipoprotein genes and the surrounding genes.

In *M. tuberculosis* and *M. bovis, lpqT* (*Rv1016c*), *lppE* (*1881c*), *lppO* (*Rv2290*), *lprE* (*Rv1252c*) and *lpqF* (*Rv3593*) were co-transcribed with their surrounding or proximal genes; which indicates that they are functionally connected with *prsA* (*Rv1017c*), *cyp140* (*Rv1880c*), *Rv2291*, Rv1251c and *Rv3592*, respectively (Fig. 1 and Table 1).

**Figure 1:**
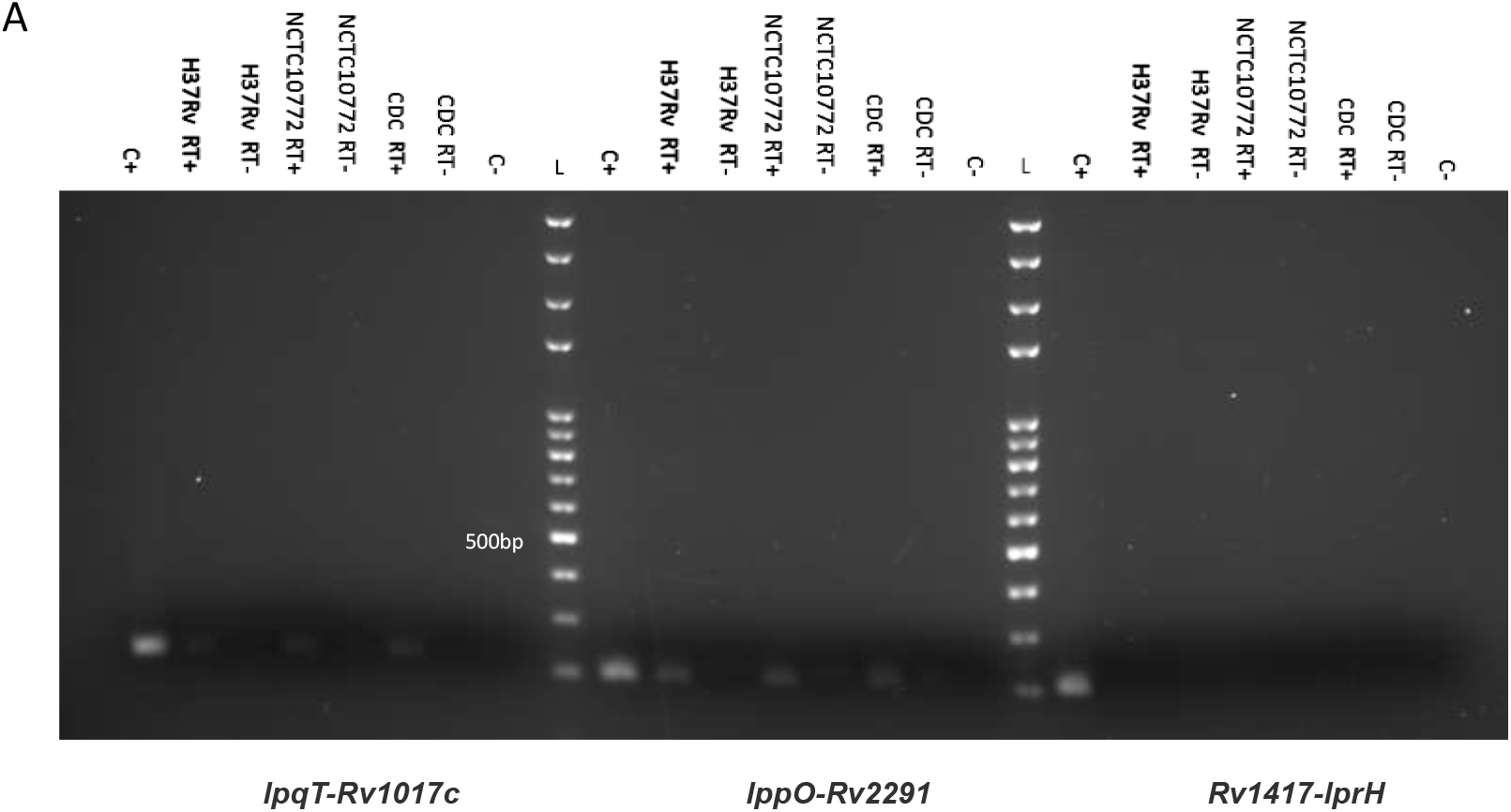
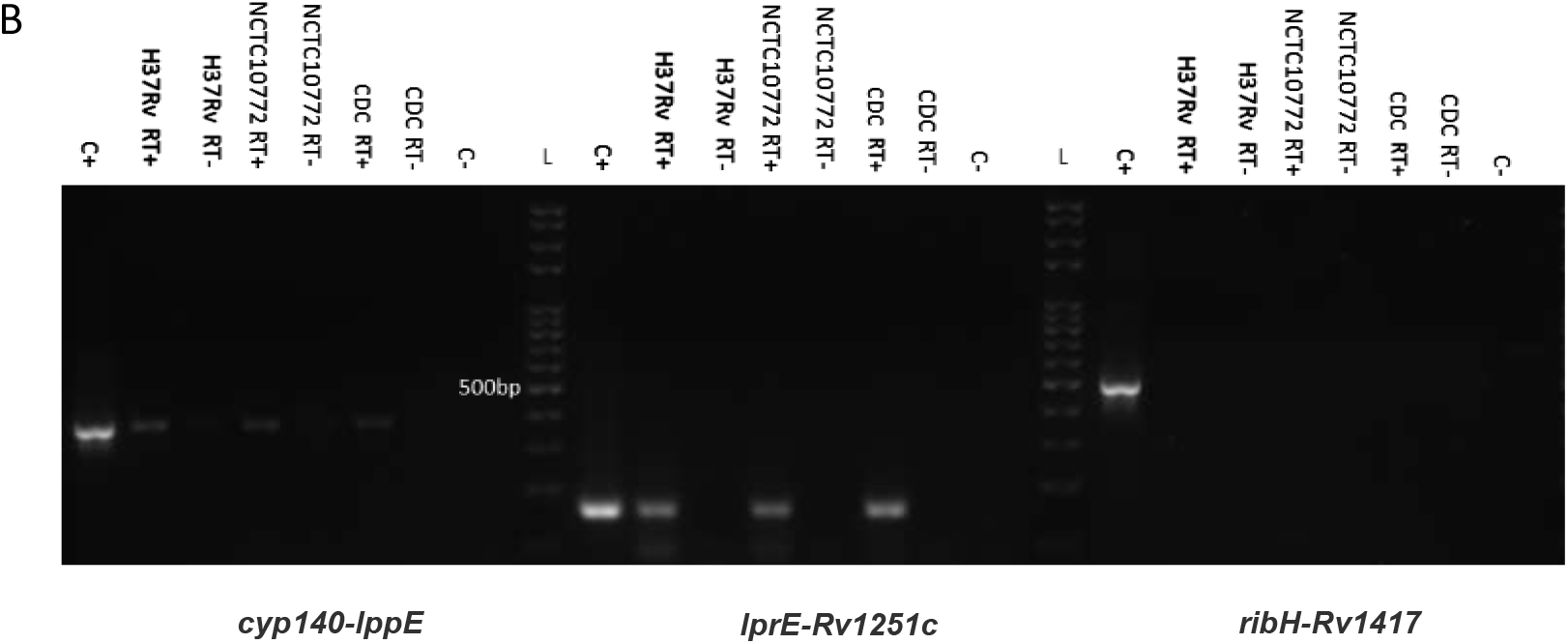
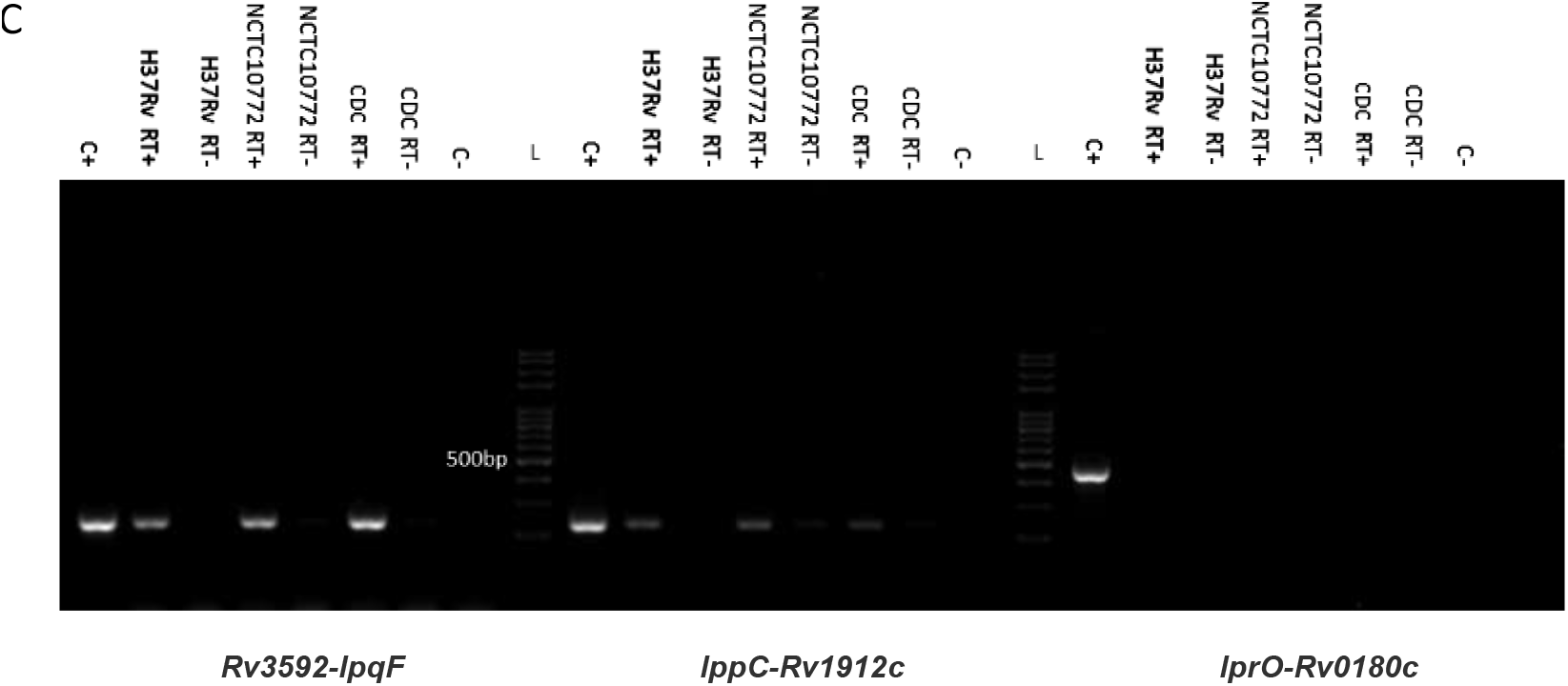
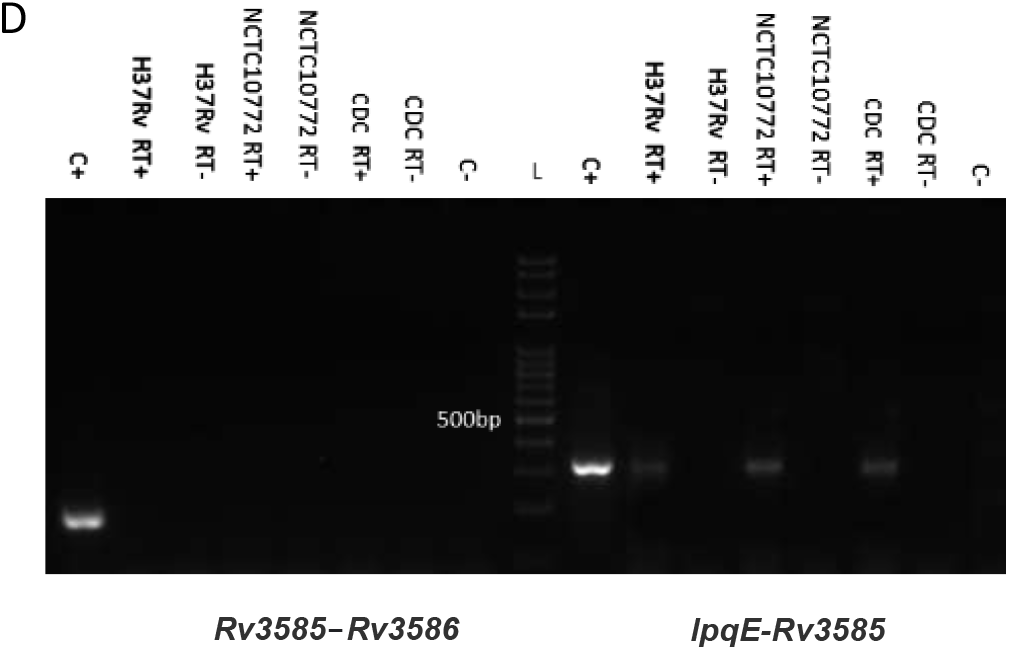
RT-PCR amplification of the different intergenic regions: *lpqT-Rv1017c*; *lppO-Rv2291*; *Rv1417-lprH* (A) *cyp140-lppE*; *lprE-Rv1251c*; *ribH-Rv1417* (B) *Rv3592-lpqF*; *lppC-Rv1912c* (C) *Rv3585-Rv3586* and *lpqE-Rv3585* (D). Purified DNA from CDC (50ng) was used as the positive control for the PCR reactions (C+) and 1uL of water as negative control (C-). The products were analysed in 2% agarose gels. The lines for the DNA ladder (100bp TransGen Biotech) were intercalated between the different intergenic regions (L).

Regarding the function of the proteins codified by the surrounding or proximal genes, PrsA is involved in the synthesis of a central metabolite required for producing arabinogalactan or lipoarabinomannan (Alderwick et al., 2011). This suggests a role for LpqT in cell wall remodelling. In addition, Cyp140 is a putative cytochrome P450 140, which is encoded in close proximity to Rv1882c —a probable short-chain type dehydrogenase/ reductase— and Rv1883c —a protein carrying a polyketide cyclases/ dehydrases domain. Therefore, this predicted operon may be essential for a cell wall biosynthesis process (Johnston et al., 2009). Rv2291 is a putative thiosulfate sulfurtransferase, suggesting a role for LppO in sulfur metabolism, whereas Rv3592 encodes a protein that participates in heme degradation (Rittershaus et al., 2018). This result may indicate that LpqF participates in iron metabolism. Finally, Rv1251c is a putative endonuclease with an ATPase domain. In this respect, the endonuclease and ATPase activities have been associated to enzymes involved in DNA translocation (Meisel et al., 1995).

The RT-qPCR experiments, however, failed to demonstrate that LprH (Rv1418) is encoded in the same transcriptional unit as Rv1417 or that *lpqE* are in the same transcriptional units as *disA* —a gene encoding an enzyme involved in the synthesis of cyclic-di-AMP, which is a second messenger (Bai et al., 2012).

## Conclusions

Based on the *in silico* analysis of the genomic structure of lipoprotein-encoding genes and RT-PCR experiments, we propose new possible roles for five lipoproteins of *M. tuberculosis*. Two of these lipoproteins are likely involved in cell wall biogenesis processes.

## Author Approvals

All authors have seen and approved the manuscript, and that it hasn’t been accepted or published elsewhere.

